# Distinct neurocomputational mechanisms support informational and normative conformity

**DOI:** 10.1101/728428

**Authors:** Ali Mahmoodi, Hamed Nili, Dan Bang, Carsten Mehring, Bahador Bahrami

**Author notes:** These authors contributed equally to this work (Dan Bang, Carsten Mehring, Bahador Bahrami).

## Abstract

A change of mind in response to social influence could be driven by *informational* conformity to increase accuracy, or by *normative* conformity to comply with social norms such as reciprocity. Disentangling the behavioural, cognitive and neurobiological underpinnings of informational and normative conformity have proven elusive. Here, participants underwent fMRI while performing a perceptual task that involved both advice-taking and advice-giving to human and computer partners. The concurrent inclusion of two different social roles and two different social partners revealed distinct behavioural and neural markers for informational and normative conformity. dACC BOLD response tracked informational conformity towards both human and computer but tracked normative conformity only when interacting with human. A network of brain areas (dmPFC and TPJ) that tracked normative conformity increased their functional coupling with the dACC when interacting with humans. These findings enable differentiating the neural mechanisms by which different types of conformity shape social changes of mind.

## Introduction

We are often faced with opinions that are different from our own. In these situations, we sometimes decide to stick to our own opinion and other times we change our mind. One key factor to select between these opposite social behaviours is our sense of confidence: the lower the confidence in our initial opinion, the higher the probability that we change our mind(1,2). However, the way in which we process differing opinions has been shown to be influenced by a range of factors some of which are unrelated to accuracy, such as a desire to fit in with a group (3) or how receptive others have previously been toward us (4). How people balance these epistemic and social factors remains an open and fundamental question in social cognitive neuroscience. Here, we develop an empirical framework for understanding the mechanisms that underpin social changes of mind at the cognitive and neural level.

There has been a recent interest in understanding changes of mind in non-social situations (1,5–7). In a typical experiment, people are given the option to change their mind about an initial decision after being presented with additional (post-decision) evidence. People have been shown to solve this problem by computing the probability that the initial decision was correct given all the evidence, and markers of neural activity obtained with fMRI have revealed that this confidence computation is supported by dorsal anterior cingulate cortex (dACC) (1). Interestingly, dACC also appears to play a central role in changes of mind in social situations (8). For example, an initial set of fMRI studies focused on situations where people changed their subjective preferences (e.g., facial-attractiveness ratings) after observing those of others (9–11). Activity in dACC was found to track the observed difference between one’s own and others’ preferences and in turn predict whether people changed their reported preferences to align with those of others (9–11). A more recent fMRI study asked people to make a perceptual decision after observing the recommendation of an advisor (12), and found that dACC activity tracked whether or not people based their decision on the advisor’s response.

Traditionally, social influence – and thereby the factors that enter into social changes of mind – has been classified as informational or normative (13). Informational influence is when we change our beliefs towards those of others in order to maximise accuracy. As in non-social situations, this process is likely to be governed by our sense of confidence in our own initial beliefs. By contrast, normative influence is when we change our beliefs towards those of others for reasons that are unrelated to accuracy. For example, we may seek to maximise group cohesion or social acceptance (3). The challenge is that, while informational and normative factors are often in direct competition, they may nevertheless drive similar behavioural responses. For example, in the fMRI studies on subjective preference, a common interpretation is that people adapted their reported preferences towards others because they felt a pressure to conform to the group. However, people may have been uncertain about their own preferences (14) and used others as a cue to infer what they themselves feel (15). Similarly, in studies where people made a perceptual decision after observing the recommendation of an advisor, people may have based their decision on the advisor’s response because they felt that it was the socially right thing to do or because they genuinely had low confidence in their own sensory percept. It therefore remains an open question how the brain balances informational and normative factors during social changes of mind.

In the current study, we investigated the cognitive and neural basis of social changes of mind, by using a social perceptual decision task that directly separates informational and normative factors. On each trial, participants first made a perceptual estimate and reported their confidence in this response and were then presented with a partner’s perceptual estimate. On one half of trials, participants had the opportunity to revise their estimate. On the other half, the partner did. Unbeknownst to participants, we manipulated the degree to which the partner’s revised estimate was influenced by the participant’s initial estimate. Critically, this manipulation and concurrent sending and receiving social influence unveils a normative reciprocity effect: participants are influenced more by the partner who has, in turn, been influenced more by them, regardless of task performance (Mahmoodi et al., 2018). In this way, one can separately measure the contribution of informational factors (the degree to which participants feel that they could improve on their perceptual estimate by taking into account that of the adviser, which is indexed by participants’ confidence in their initial estimate) and normative factors (the degree to which participants feel that they should reciprocate influence, which is controlled by our manipulation of partner behaviour) to social changes of mind.

To anticipate our results, behavioural analyses revealed that participants’ revision of their perceptual estimate was governed by both informational and normative factors: they shifted more towards the partner’s estimate when they had low confidence in their own estimate and when there was a higher demand for reciprocating influence. Critically, in a control condition, where participants were told that they interacted with a computer, changes of mind were only influenced by informational factors. Analysis of fMRI data showed that dACC activity tracked both confidence and the demand for reciprocating influence at the time of revision. In line with the behavioural results, dACC activity tracked the normative factor only when participants believed that they worked with a human (but not a computer) partner. Further, we found that traditionally social brain areas – dorsomedial prefrontal cortex (dmPFC) and temporoparietal junction (TPJ) – tracked the degree to which a partner took into account the participants’ perceptual estimate on trials where the partner revised their estimate and in turn increased its coupling with dACC on trials where the participant revised their estimate when both informational and normative demands were high. Taken together, these results support a general role for dACC in coordinating changes of mind in both non-social and social situations.

## Results

### Experimental task

Participants (N = 60) performed a social perceptual decision-making task. In each experimental session, three participants came to the lab at the same time. The three participants met briefly and had their individual photos taken by the experimenter. One of the three participants performed the task while undergoing fMRI (N = 20) and the remaining two participants performed task in separate behavioural testing booths (N = 40). The task consisted of four blocks of trials (scan runs) – with each participant paired with a unique partner in each block. Participants were told that in two of the four blocks, the partner was a computer. In each of the other two blocks, the participant was paired with one of the other participants. In reality, unbeknownst to the participants, all four partners were simulated. To help participants separate the partners, and to strengthen the computer-human distinction, participants were shown the photo of the current partner at the beginning of each trial.

In each trial, participants privately made a perceptual estimate about the location of a visual target and then rated their confidence in this estimate on a scale from 1 to 6 (Figure 1A). After having indicated their estimate and confidence, participants saw the partner’s estimate of the location of the same visual target. The partner’s estimate was generated by drawing a random sample from a Von Mises distribution centred on the correct answer. On odd (observation) trials, participants waited while the partner revised their estimate in light of the participants’ estimate and were then shown the partner’s revised estimate. On even (revision) trials, participants had the opportunity to revise their estimate in light of the partner’s estimate, after which the revised estimate was shared with the partner. To ensure that participants paid attention to the partner’s estimate, they were required to place their revised estimate between their initial estimate and that of the partner. Participants did not receive feedback.

**Figure 1:**
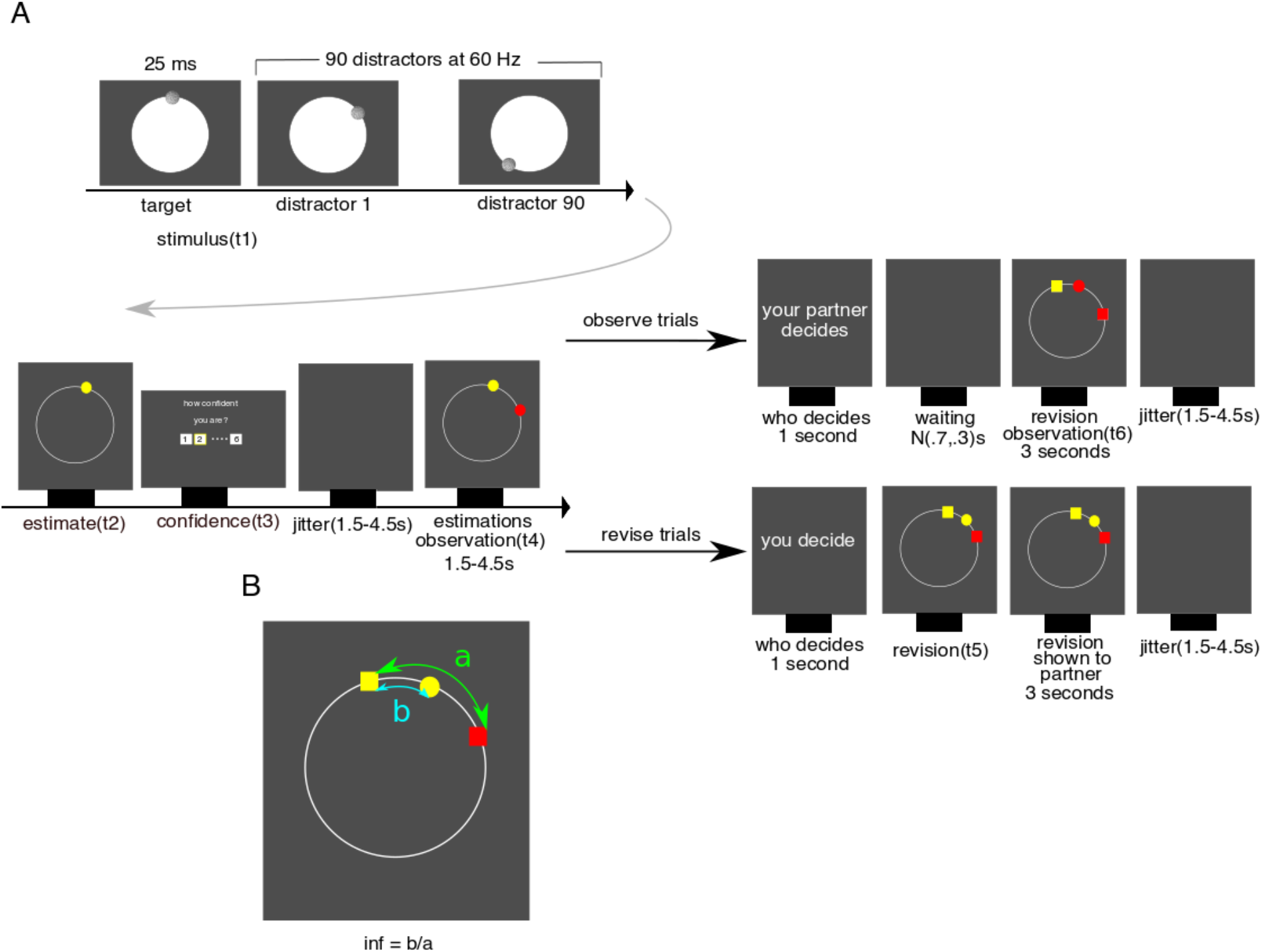
Multi-stage decision-making framework for studying social changes of mind. **A** On each trial, participants observed a sequentially presented series of dots on the screen (t1). They were then required to indicate where the very first dot in the series appeared (yellow dot; t2) and report their confidence on a discrete scale from 1 to 6 (t3). Next, they were presented with the estimate of a partner (red dot; t4) concerning the same stimulus. On half of the trials (observe), the partner had the opportunity to change their mind, and participants only observed the partner’s revision. On the other half of trials (revise), participants had the opportunity to revise their initial estimate (t5). Participants were informed whether they were paired with computer or a human in each block. **B** angular distance between initial estimates = a; angular distance between estimates after revision = b; social influence = b/a.

We manipulated the influence that participants exerted over their partners on observation (odd) trials: one human and one computer partner were strongly influenced by participants’ estimate (susceptible blocks), whereas one human and one computer partner were only slightly influenced by participants’ estimate (insusceptible blocks). In this way, the observation (odd) trials allowed us to introduce a normative aspect to the task – the degree to which the partner shifted towards the estimate made by the participant – whereas the revision (even) trials allowed us to quantity the impact of informational (confidence) and normative (influence) factors on social changes of mind.

### Behavioural separation of informational and normative factors

To disentangle the contribution of informational and normative factors to social changes of mind, we performed a linear mixed-effects regression analysis. We predicted that, in revision trials, the degree to which participants changed their estimate towards that of the partner (from here onwards, *revision*) depends on the participant’s confidence in their initial estimate, the influence that they exerted over their partner on the previous trial (i.e., *influence*) and the interaction between confidence and influence. In the interest of clarity, we define revision and influence separately here (see Figure 1B). Revision is defined by the angular difference between participant’s initial and final estimates divided by the angular difference between the two initial estimates. Influence is defined by the angular difference between the partner’s initial and final estimates divided by the angular difference between their two initial estimates. Informational conformity would be demonstrated by a negative correlation between confidence and revision recorded in the same trial. A positive correlation between revision in the current trial and influence in the previous trial would be evidence for normative conformity. Critically, we would only expect this latter relationship to be observed for the human partner. This is because normative influence should, by definition, not pertain to human-computer interactions (4). Therefore, for the human condition we predict a posotove effect of influence on revision and a potential negative effect the interaction between confidence and influence as these the effect of one might depend on the other.

Our first model (LLM1) also included a term for the partner type (human or computer) and interactions between partner type and our three variables of interest. In support of the rationale behind our task design, we found that confidence had a negative effect on revision (parameter estimate: −0.22, 95% CI: [−.42 −.02], t(3532) = −2.16, p = .03), whereas influence had a positive effect (parameter estimate 0.34, 95% CI [.07 .26], t(3532)= 2.49, p= .01). Critically, there was also an interaction between partner type and influence (parameter estimate: −0.18, 95% CI: [−.35 −.01], t(3532) = 2.08, p = .03) – indicating that the effect of influence was different for human and computer partners.

To unpack these results, we ran separate models for each partner type (LMM2). For the computer partner, there was a negative effect of confidence on revision (parameter estimate: −.37, 95% CI: [−.48 −.26], t(1766) = −6.51, p<.001) but no effect of influence (parameter estimate: 0.01, 95% CI: [−.1 .14], t(1766) = .29, p= .77) and no interaction between confidence and influence (parameter estimate −0.03, 95% CI [−.21 .14], t(1766) = −.37, p= .7) (Figure 2). For the human partner, confidence had a negative effect on revision (parameter estimate −.3, 95% CI [−.41 −.2], t(1766) = −6.02, p<.001) and – in line with normative concerns being specific to human-human interactions – there was a positive effect of influence (parameter estimate 0.17, 95% CI [.05 .28], t(1766)= 2.89, p= .003) and a negative effect interaction between confidence and influence (parameter estimate −0.21, 95% CI [−.4 −.03], t(1766) = −2.35, p= .01) (Figure 2) – in other words, the higher the confidence, the lower the effect of influence on revision, and vice versa.

**Figure 2:**
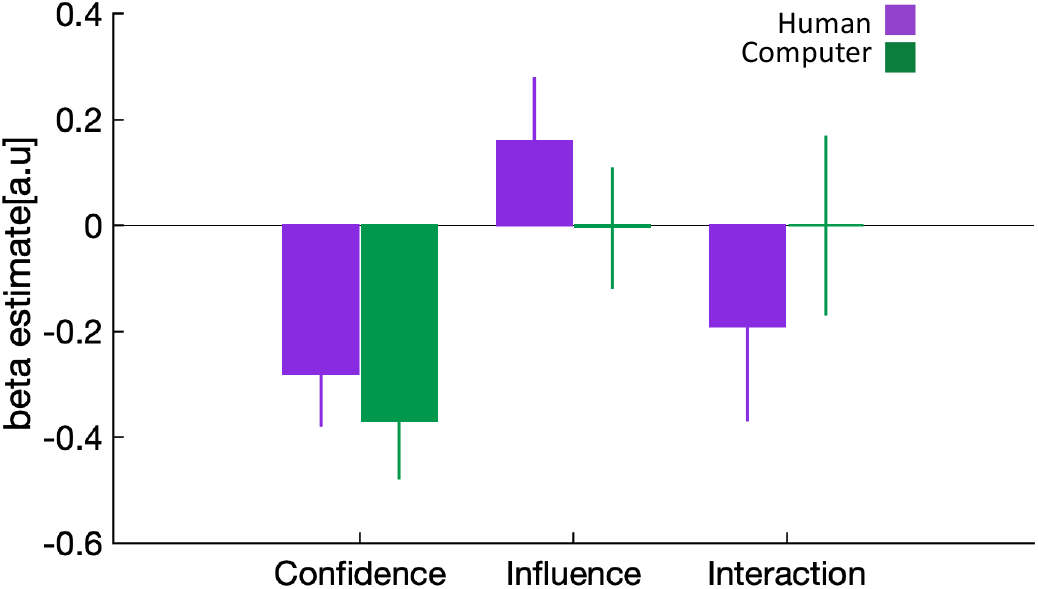
Behavioural separation of informational and normative factors. We ran linear mixed-effects models separately for human and computer conditions in which we predicted participants’ revision toward their partner using confidence, influence and their interaction. Data are represented as group mean ± 95% confidence intervals.

And finally, the average revision value was .34, indicating that participants overall gave more weight to their own initial estimate (Wilcoxon sign-rank test against .5, median = .35, W = 582, p < .0001) – consistent with the finding that, all else being equal, people tend to discount the opinion of others (16).

### Encoding of informational and normative factors in dACC

Having established behaviourally that our experimental task dissociated the influence of informational and normative factors on social changes of mind, we next turned to the fMRI data to identify neural substrates that may support the integration of these factors into a social change of mind. We focused our analyses on dACC as this area has consistently been linked to changes of mind in social (8) and non-social situations (1). Our key question was whether dACC tracks both informational and normative factors, or only one of these factors, during social changes of mind. We used the same dACC ROI as in the study by Fleming et al. (2018) while noting that this ROI overlaps with the dACC clusters identified in the social studies discussed in the Introduction.

We first asked whether dACC encodes participants’ reported confidence at the time of the initial estimate (t2 in Figure 1). As shown by the behavioural results, confidence is a central component of informational conformity as it indexes the degree to which participants feel that they can improve on their perceptual estimate by taking into account that of a partner – regardless of whether the partner is human or a computer. GLM analysis of activity time courses locked to the onset of the estimation screen showed that dACC tracked confidence negatively on both human and computer trials (aggregated across both condition, Wilcoxon sign-rank test, p = .002, W = 24), (Figure 3) – a result which is consistent with the literature on the neural basis of confidence in non-social and social settings (17,18).

**Figure 3:**
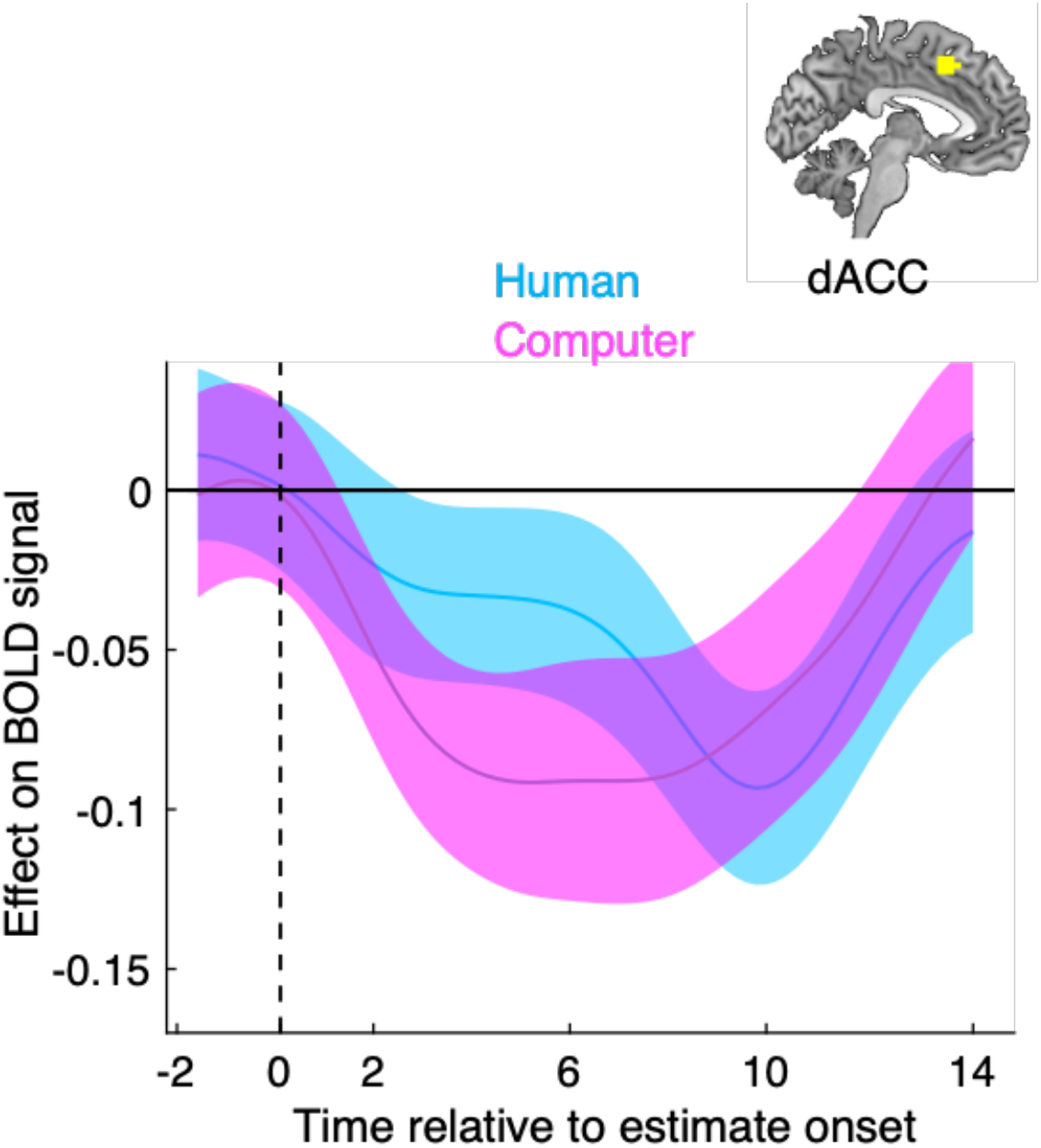
Validation of the reported relationship between dACC activity and decision confidence. GLM analysis of the effect of reported confidence on dACC activity time courses locked to the onset of the perceptual estimation screen. The group level significance was estimated using a leave-one-out procedure. Data are represented as group mean ± SEM. Vertical dashed line indicates estimation onset (t2).

We next asked whether dACC encodes the normative component of social changes of mind – the demand to reciprocate social influence – when the participant revises their initial estimate. Additionally, we asked whether dACC continues to encode decision confidence at this stage. To this end, we performed a GLM analysis of activity time courses locked to the onset of the revision screen (t5 in Figure 1) using confidence, influence and their interaction as predictors. Because the behavioural results showed that influence (its main effect or interaction with confidence) did not affect revision in the computer condition, we performed this analysis separately for the human and computer conditions. In line with the previous results, dACC tracked confidence at the time of revision in both the human and computer conditions aggregated across both condition (aggregated across conditions, Wilcoxon sign-rank test, p = .01, W = 170), (Figure 4). Interestingly, dACC also tracked the interaction between confidence and influence at the time of revision but, critically, only in the human condition (Wilcoxon sign-rank test, p = .02, W = 46), (Figure 4). The interaction effect means that in the human condition, the response of the dACC to confidence was lower if influence was high and vice versa. Notably, we reached the same conclusion when we repeated this analysis using an alternative model (see Supplementary Material).

**Figure 4:**
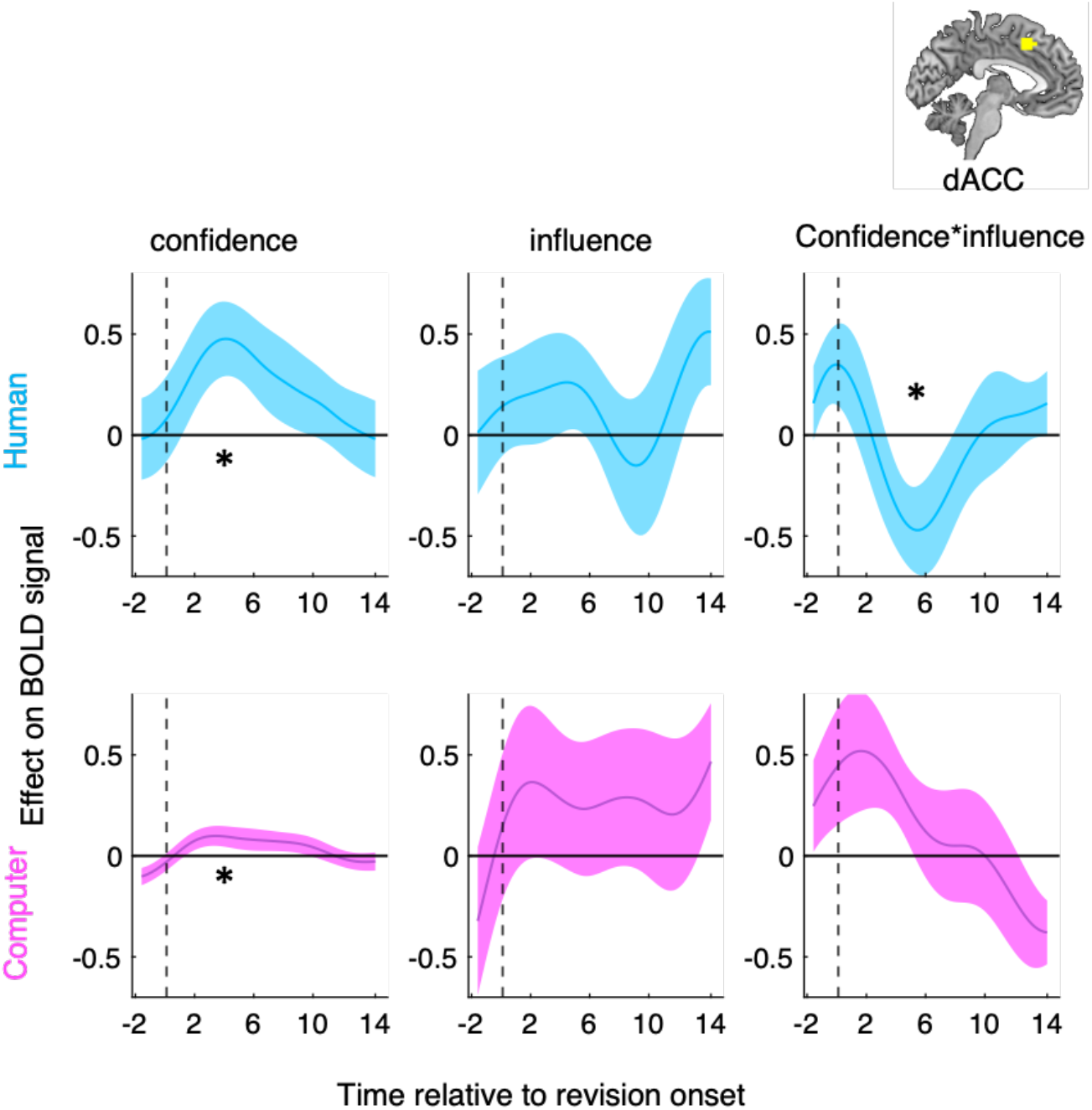
dACC tracks informational and normative factors during social changes of mind. GLM analysis of the effects of reported confidence, social influence and their interaction on dACC activity time courses locked to the onset of the revision screen for the human (top) and the computer (bottom) condition. The star indicates that the time course was significantly different from zero using a leave-one-out procedure. Data are represented as group mean ± SEM. Vertical dashed line indicates revision onset (t5).

Next, we focused on the human condition and used the division of trials into revision and observation trials (Figure 1) to test whether the hypothesis that dACC tracked informational and normative factors specifically in the service of social changes of mind. If the dACC response pattern is driven by the prospect of having to make a second estimate, then we expect to find the encoding of informational and normative factors only in revision and not in observation trials. If, on the other hand, the task variables were automatically encoded regardless of current task requirements, then we would expect to see the dACC response in both trial types. To test this prediction, we used dACC activity time courses locked to the onset of the screen that announced whether the current trial was a revision or an observation trial – notably, the screen appeared after participants had made their initial estimate and had been presented with that of their partner. In support of a specific role of dACC in revision, this analysis revealed that dACC did not track confidence, influence or their interaction on observation trials (Figure S1).

### Encoding of normative factors in social brain areas

Having established that dACC integrates informational and normative factors during social changes of mind, we sought to identify the neural substrates that were most likely to provide the normative input to dACC. The informational component, i.e., confidence, is immediately available on a revision trial, and may be encoded by dACC itself (1). The normative component, i.e., social influence, however, must first be assessed on the preceding observation trial and then carried forward to the upcoming revision. We hypothesised that dorsomedial prefrontal cortex (dmPFC) and the temporoparietal junction (TPJ) may serve such assessor function. Both of these areas are part of the theory of mind network(19–21) and have been shown to track the trial-by-trial variation in task-relevant social variables(22,23) and social prediction(24). To test this hypothesis, we first focused on observation trials and performed a GLM analysis of dmPFC and TPJ activity time courses locked to the observation of partner’s change of mind (t6 in Figure 1) using the influence as a predictor. In line with our hypothesis, both dmPFC (Wilcoxon sign-rank test, p = .001, W = 189), and TPJ (Wilcoxon sign-rank test, p = .01, W = 173), tracked influence at the time of revision observation in the human condition (Figure 5A–B).

**Figure 5:**
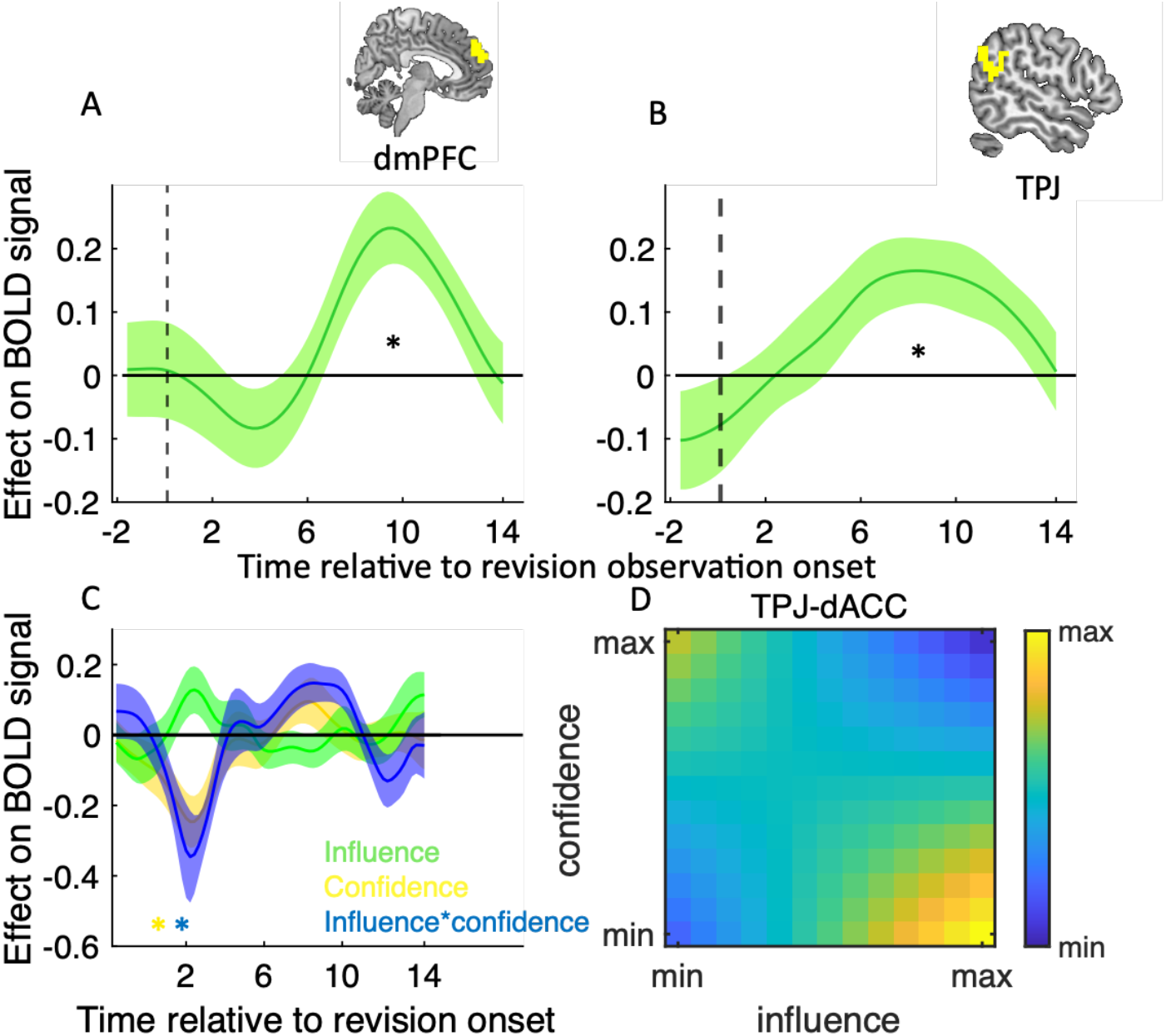
Encoding of normative factors in social brain areas. **A,B** dmPFC and TPJ tracks social influence on observation trials. GLM analysis of the effects of social influence on (left panel) dmPFC and (right panel) TPJ activity time courses locked to the onset of the revision observation screen. **C** Psychophysiological interaction analysis of ROI activity time courses. Traces are coefficients from a GLM in which we predicted dACC activity from the interaction between TPJ activity and (1) confidence (yellow), (2) influence (green) and (3) the interaction between confidence and influence (blue) – while controlling for the main effect of each term (confidence, influence and TPJ activity). ***D*** Visualisation of TPJ-dACC connectivity. Hotter colours indicate greater TPJ-dACC connectivity as a function of variation (in *z*-score units) in influence (x-axis) and confidence (y-axis). TPJ-dACC connectivity was estimated using group-level coefficients averaged across a time window from 2 s to 3 s. In **A-B,** data are represented as group mean ± SEM. In A, B, anc *C,* the star indicates that the time course was significantly different from zero using a leave-one-out procedure.

If dmPFC and/or TPJ provide the normative factor that is used by dACC to drive social changes of mind, then connectivity between these areas and dACC on revision trials should vary with the demand for reciprocating influence as computed on a preceding observation trial. To test this prediction, we performed a psychophysiological interaction (PPI) analysis in which we quantified connectivity between dmPFC/TPJ and dACC at the onset of the revision screen on human trials. This was done using a GLM where we predicted the activity of the dACC as a function of confidence, influence, TPJ activity and all interaction terms. We included confidence, influence and its interaction with influence as the behavioural factors as our behavioural and neural results from the dACC show that the contribution of influence to social changes of mind depends on confidence, and vice versa (Figure 2 and Figure 4). In support of our hypothesis, this analysis revealed a close coupling between TPJ and dACC. Shortly after the onset of the revision screen, TPJ-dACC connectivity varied with confidence (Wilcoxon sign-rank test, p = .007, W = 33), and its interaction with influence (Wilcoxon sign-rank test, p = .002, W = 25), (Figure 5C). Visualisation of these effects showed that TPJ-dACC connectivity was highest when influence was high and confidence was low (Figure 5D) – the condition where normative factors have the largest influence on social changes of mind (Figure 2). There was also a peak when confidence was high and influence was low. We note that dmPFC-dACC connectivity showed the same pattern as TPJ-dACC connectivity but did not reach significance (Figure S2).

Finally, to complement our ROI analysis, we performed an exploratory whole-brain analysis in which we searched for neural correlates of our variables of interest. We confirm that we found the neural correlates of confidence but not influence and the interaction between confidence and influence at the whole brain level (See GLM1 in Methods and supplementary Material for the full details of the whole brain analysis).

## Discussion

A key feature of adaptive behavioural control is our ability to change our mind as new evidence comes to light. Previous research has identified dACC as a neural substrate for changes of mind in both non-social situations, such as when receiving additional evidence pertaining to a previously made decision (1), and social situations, such as when weighing up one’s own decision against the recommendation of an advisor (12). However, unlike the non-social case, the role of dACC in social changes of mind can be driven by different, and often competing, factors that are specific to the social nature of the interaction (13). In particular, a social change of mind may be driven by a desire to be correct, i.e., informational influence. Alternatively, a social change of mind may be driven by reasons that are unrelated to accuracy – such as social acceptance – a process called normative influence. To date, studies on the neural basis of social changes of mind have not disentangled these processes. It is therefore unclear how the brain tracks and combines informational and normative factors (12,25,26).

Here, we leveraged a recently developed experimental framework that separates humans’ trial by trial conformity into informational and normative components (4) to unpack the neural basis of social changes of mind. On each trial, participants first made a perceptual estimate and reported their confidence in this response. In support of our task rationale, we found that, while participants’ changes of mind were affected by confidence (informational) in both human and computer settings, they were only affected by the need to reciprocate influence (normative) in the human setting. It should be noted that participants’ perception of their partners’ accuracy is also an important factor in social change of mind (we tend to change our mind toward the more accurate participants). Not being the focus of this study, we therefore controlled for the effect of partners’ accuracy by designing partners of equal performance. Our previous study showed that first, participants’ perceived performance of their partner was not different across conditions, and second the effect of confidence and influence on social change of mind could not be explained away by their perception of their partners’ accuracy(4).

Building on previous research on the neural basis of changes of mind (1,10,12), our analysis of fMRI data acquired during task behaviour focused on dACC, and in particular the degree to which dACC encoded informational and normative factors in the different conditions of our task design. Overall, our neural results support a central role for dACC in orchestrating social changes of mind. First, in line with the behavioural results, when participants were given the opportunity to revise their initial estimate, dACC tracked confidence only in the computer setting but confidence as well as the need to reciprocate influence in the human setting. Finally, in the human setting, dACC tracked confidence and the need to reciprocate influence only when participants had the opportunity to revise their initial estimate (revision trials) and not when it was the partner’s turn to do so (observation trials) – demonstrating that the dACC responses were directly tied to behavioural control. More broadly, looking beyond changes of mind, our neural results are in line with a proposal that all task-relevant variables, independent of their origin and nature, converge in dACC and that dACC in turn supports the selection of task-appropriate behavioural responses (27).

If dACC supports the integration of informational and normative factors into a social change of mind, which brain regions provide the respective inputs to this process? As for the informational component of a social change of mind, our results suggest, at first glance, that dACC itself may be involved in the construction of decision confidence. In particular, we found that, at the time of making a perceptual estimate, dACC tracked participants’ confidence in this estimate – a temporal association identified by other studies on the neural basis of decision confidence (18). However, recent research, which disentangled the components of decision confidence (28) or separated a sense of confidence from explicit confidence reports (17), suggest that this temporal association is due to a role of dACC in controlling confidence-based behaviours rather than encoding a sense of confidence per se. However, we acknowledge that our whole-brain analysis did not reveal any other brain regions that may have supported a confidence computation. As for the normative component of a social change of mind, our analysis of TPJ and dmPFC – both parts of the so-called theory of mind network (19,20) – suggests that these regions may provide the social context for a change of mind. First, on observation trials – when observing a human partner’s revised estimate – TPJ and dmPFC tracked the degree to which a partner took into account the participant’s estimate and thereby the degree to which participants should reciprocate influence on the subsequent revision trial. Second, on revision trials – when required to balance informational and normative considerations – functional coupling between dACC and TPJ was highest when the normative component of a social change of mind was required in the conformity process.

Our results may prompt a reconsideration of earlier accounts of the role of dACC in social changes of mind. For example, Qi et al. (2018) found that dACC activity predicted the degree to which participants’ perceptual judgements deviated from those of an advisor (12). Invoking the conflict monitoring theory of dACC function (29), Qi et al. (2018) took this response pattern to suggest that dACC tracks social conflict. However, as highlighted by our study, decision confidence and social conflict are often two sides of the same coin – the higher the degree of confidence, the lower the influence of others – making it hard to arbitrate between a conflict account of dACC and its role in encoding informational and normative components of conformity. In order to test the social conflict account of dACC, we reran the GLM analysis of dACC activity time courses locked to the onset of the revision screen, including the difference between participants’ initial estimate and the partner’s estimate as well as the difference between participants’ revised estimate and the partner’s estimate in addition to confidence, the need to reciprocate and the interaction between confidence and the need to reciprocate. Notably, neither of the difference terms – both markers of social conflict as quantified by Qi et al. (2018) – were encoded by dACC. While our results do not rule out that dACC may track social conflict, they show that social conflict does not provide a unified explanation of dACC function during social changes of mind. Rather, dACC appears to track any variable – irrespective of whether it is informational or normative in nature – that are deemed relevant in the context of the current task at hand.

Our results showed that humans can and do interact with non-human social partners via informational but not normative conformity. When humans interact with an inanimate computer partner, normative conformity is not observed neither in behaviour nor in human brain. This will have important ramifications for the new and burgeoning field of human-AI interactions. For example, with the imminent introduction of self-driving cars into everyday life, studies such as ours will be able to help anticipate the emergence of norms of politeness between human and AI drivers on the road

In summary, our results suggest the cooccurrence of informational and normative conformity in the humans’ social change of mind but only when working with other humans. Our results also suggest a broader role for the dACC in change of mind in both social and non-social contexts. Our results imply that the social brain areas (TPJ/dmPFC) might make a network together with the dACC to include the normative components of social influence in the conformity process.

## METHODS

### Participants

In total, 60 healthy adult participants (30 females, mean age ± std:25 ± 3) participated in the experiment after having given written informed consent. The experimental procedure was approved by the ethics committee at the University College London (UCL).

### Experimental paradigm

Participants were presented with a sequence of 91 visual stimuli consisting of small circular Gaussian blobs (r = 5mm) in rapid serial visual presentation on the screen. The first stimulus was presented for 30ms while every other stimulus was presented for 15ms each. Participants’ task was to identify the location of the first stimulus. Participants were required to wait until the presentation of all stimuli were finished, and then indicate the location of the first stimulus using a keyboard. The reported location was marked by a yellow dot. After participants reported their initial estimate, they were required to report their confidence about their estimate on a numerical scale from 1 (low confidence) to 6 (high confidence). For participants in the fMRI scanner, this stage was followed by a blank jitter randomly drawn from a uniform distribution from 1.5-4.5 seconds. Afterwards, participants were shown the estimate of their partners about the same stimulus for 1.5 seconds by a small red dot on the screen (plus a jitter time randomly drown from a uniform distribution from 1.5-4.5 seconds for the fMRI experiment). Then, either the participant revised her estimate or observed the partner revise theirs. After the second estimate was made, all estimates were presented to the participants for 3 seconds (plus a jitter time randomly drawn from a uniform distribution from 1.5-4.5 seconds for the fMRI experiment). In this stage, the first estimate was shown by a hexagon (for participants outside the fMRI scanner) or by a dot with a different colour (for the fMRI experiment) to be distinguished from the second estimate which was shown by a circle (Figure 1 B). Participants were told that their payoff would be calculated based on the accuracy of their first and second estimates. However, everyone was given a fixed amount at the end of the experiment. In the fMRI experiment, 10 participants’ dot colour was yellow and their partner’s dot colour was red. For the remaining 10 participants, the colours were reversed. Further details of the experimental paradigm are described in our previous study (4).

**T**hree participants came to the MRI facilities at the same time. After reading the task instructions, one participant was selected to perform the experiments in the fMRI scanner while the other two carried out the behavioural task outside the scanner. Participants were told that they will play with four different partners: two human partners (the two they met before the experiment) and two computer partners which were controlled by the algorithm described below. Participants completed 4 blocks (scan runs) of the experiment each consisting of 30 trials. In each block they only worked with one partner. At the beginning of each trial, a photo of the partner they work with was shown to the participants. Photos of two different computers with different colours (counterbalanced across participants) represented the two computer partners. In reality, and unknown to the participants, all partners’ estimates were generated by a computer algorithm. The partners only differed in the way they generated their second choice. In the insusceptible blocks, participants’ influence over their partner was chosen randomly from a uniform distribution on the interval [0, 0.2]. For the susceptible partner, participants’ influence was chosen with a probability of 0.5 from a uniform distribution on the interval [0.7, 1], with a probability of 0.2 from a uniform distribution on the interval [0.3, 0.7], and with a probability of 0.3 from a uniform distribution on the interval [0, 0.3].

All experiments were performed using Psychophysics Toolbox (30) implemented in MATLAB (Mathworks). The behavioural data were analysed using MATLAB.

### Debriefing

After each session of the experiment, all participants were debriefed to assess to what extent they believed the cover story. We interviewed them with indirect questions about the cover story and all participants stated that they believed they were working with other human participants in neighbouring experimental rooms (if they were told that their partner is a human partner).

### Constructing computer partners

The estimates of computer partners were calculated similarly to our previous study (4). In each trial we drew the first choice of the computer partner from a von Mises distribution centred on the target with a concentration parameter kappa=7.4 except in high confidence trials (confidence level of 5 or 6) where the partner’s choice was randomly drawn from a uniform distribution centred on the participants’ choice with a width of +/−20 degrees in the behavioural experiment and +/–50 degrees in the neuroimaging experiment.

### Linear mixed effect model 1 (LMM1)

To investigate the difference between the experimental conditions on the effect of informational and normative factors on revision we designed a model as follow:

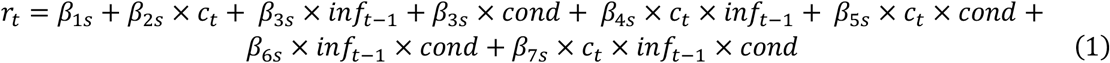

*r*_*t*_ and *c*_*t*_ correspond to the participants’ revision and confidence on trial t, respectively. *inf*_*t*–1_ corresponds to the influence that participants exerted on their partner on trial t-1. *Cond* corresponds to condition included as dummy variable (1 = human, 2 = computer). The intercept (*β*_1*s*_) and all slopes were allowed to vary across participants by including random effects of the form *β*_*ks*_ = *β*_*k*0_ + *b*_*ks*_ where 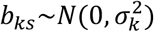.

The statistics for confidence, influence and the interaction between influence and condition are reported in the main text. The statistics for the remaining regressors were as follow: condition (parameter estimate 0.7, 95% CI [−.004 .16], t(3532)= 1.84, p= .06), confidence and influence interaction (parameter estimate −0.4, 95% CI [−.82 .01], t(3532)= −1.87, p= .06), confidence and condition interaction (parameter estimate −0.1, 95% CI [−.22 .01], t(3532)= −1.76, p= .07), and the triple interaction between confidence, influence and condition (parameter estimate 0.21, 95% CI [−.05 .47], t(3532)= 1.56, p= .11).

### Linear mixed effect model 2 (LMM2)

To investigate the distinct effect of informational and normative factors on revision we designed a model as follow:

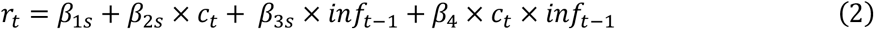

*r*_*t*_ and *c*_*t*_ correspond to the participants’ revision and confidence on trial t, respectively. *inf*_*t*–1_ corresponds to the influence that participants exerted on their partner on trial t-1. The intercept (*β*_1*s*_) and all slopes (*β*_2*s*_, *β*_3*s*_) were allowed to vary across participants by including random effects of the form *β*_*ks*_ = *β*_*k*0_ + *b*_*ks*_ where 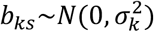. This model was fitted separately to the data from human and computer conditions. *β*_4_ was fixed across participants as the full model (where *β*_4_ varies across participants was overparametrized and the Hessian matrix was not positive definite. We therefore defined *β*_4_ as fixed effect. We note that this model has lower BIC (BIC = 745) than the models with confidence as fixed effect (BIC = 756) or influence as fixed effect (BIC = 753).

Notably, in both above models (LMM1 and LMM2) by dividing all confidence values by 6, we normalised confidence to lie between 0 and 1.

### MRI data acquisition

Structural and functional MRI data were obtained using a Siemens Avanto 1.5 T scanner equipped with a 32-channel head coil at the Birkbeck-UCL Centre for Neuroimaging. The echoplanar image) sequence was acquired in an ascending manner, at an oblique angle (≈ 30°) to the AC–PC line to decrease the impact of a susceptibility artefact in the orbitofrontal cortex with the following acquisition parameters: 44 volumes of 2 mm slices, 1 mm slice gap; echo time = 50 ms; repetition time = 3,740 ms; flip angle = 90°; field of view = 192 mm; matrix size = 64 × 64. A structural image was obtained for each participant using MP-RAGE (TR = 2730 ms, TE = 3.57 ms, voxel size = 1 mm^3^, 176 slices).

### fMRI data analysis

Imaging data were analysed using Matlab (R2016b) and Statistical Parametric Mapping software (SPM12; Wellcome Trust Centre for Neuroimaging, London, UK). Images were corrected for field inhomogeneity and corrected for head motion. They were subsequently realigned, coregistered, normalized to the Montreal Neurological Institute template, spatially smoothed (8 mm FWHM Gaussian kernel), and high filtered (128 s) following SPM12 standard preprocessing procedures.

The design matrix for GLM1 included 6 events. These were the times of stimulus representation (t1), making the first (private) estimate (t2), reporting the confidence (t3), showing the first estimates (t4), making the second (revised) estimate (t5), revision observation (t6). Furthermore, regressors t2, and t3 were parametrically modulated by subject’s reported confidence. The regressor for t5, included the parametric modulators confidence, angular distance between the participant’s own and the partner’s first estimate, angular distance between participant’s second and the partner’s first estimate, the amount of revision that participants made toward their partner’s estimate, participants’ influence over their partner in the previous trial, and the interaction between this influence and confidence. The regressor for t6, included the participants’ influence over their partner as parametric modulator. For events in which the duration depended on the participants’ reaction time (t2, t3 and t5), the natural logarithm of the reaction time i.e. log(RT) was included as the parametric modulator. Parametric modulators were not orthogonalized to allow the regressors to compete for explaining the variance.

### Regions of interest analysis

We focused on three a priori ROIs highlighted by previous research on social cognition. The TPJ mask was defined using the Human TPJ parcellation study developed by (31) and mirrored to the left hemisphere to create a bilateral mask. We used the dmPFC mask from (32) and the dACC mask from (1). We transformed each ROI mask from MNI to native space and extracted preprocessed BOLD time courses as the average of voxels within the mask. For each scan run, we regressed out variation due to head motion, and upsampled the BOLD time course by a resolution of .2 seconds. For each trial, we extracted activity estimates in a 15 s window (75 time points), time-locked to 1s before the onset of each event of interest. We used linear regression to predict the ROI activity time courses. More specifically, we applied a linear regression to each time point and then, by concatenating beta-weights across time points, created a beta-weight time course for each predictor of a regression model. We performed this step separately for each subject and pooled beta-weight time courses across subjects for visualisation. For example, for the dACC time course analysis for Figure 4, we used the following linear regression model:

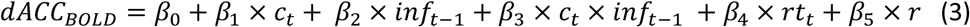

Where *c*_*t*_ correspond to the participants’ confidence on trial t and *inf*_*t*–1_ corresponds to the influence that participants exerted on their partner on trial t-1. *rt*_*t*_ corresponds to participants’ reaction time in trial t, and *r* corresponds to block number.

We tested group-level significance using a leave-one-out procedure to avoid any selection bias. For each participant and for each time course signal, we computed the peak the signal (positive or negative) for the group and calculated the beta weight of the left-out participant at the time of the group peak. We repeated this procedure for each participant and compared the resulting beta weights against zero.

## Authors Contributions

A.M., H.N., C.M., and B.B., designed the experiments. A.M. and B.B collected the neuroimaging data.

A.M. carried out the data analysis. A.M., H.N., D.B., C.M., and B.B interpreted the results. A.M. drafted the manuscript and all authors contributed to the final manuscript.

## Acknowledgments

A.M. was supported by a PhD scholarship from the Graduate School Scholarship Program of the German Academic Exchange Service (DAAD). D.B. was supported by a Sir Henry Wellcome Postdoctoral Fellowship funded by the Wellcome Trust (213630/Z/18/Z). B.B. was supported by the Humboldt Foundation, the NOMIS Foundation, and the European Research Council under the European Union’s Horizon 2020 research and innovation programme (grant agreement No. 819040 - acronym: rid-O).

## Data availability

All supporting data and code will be provided upon reasonable request to the corresponding author.

## Supplementary Material

**Figure S5:**
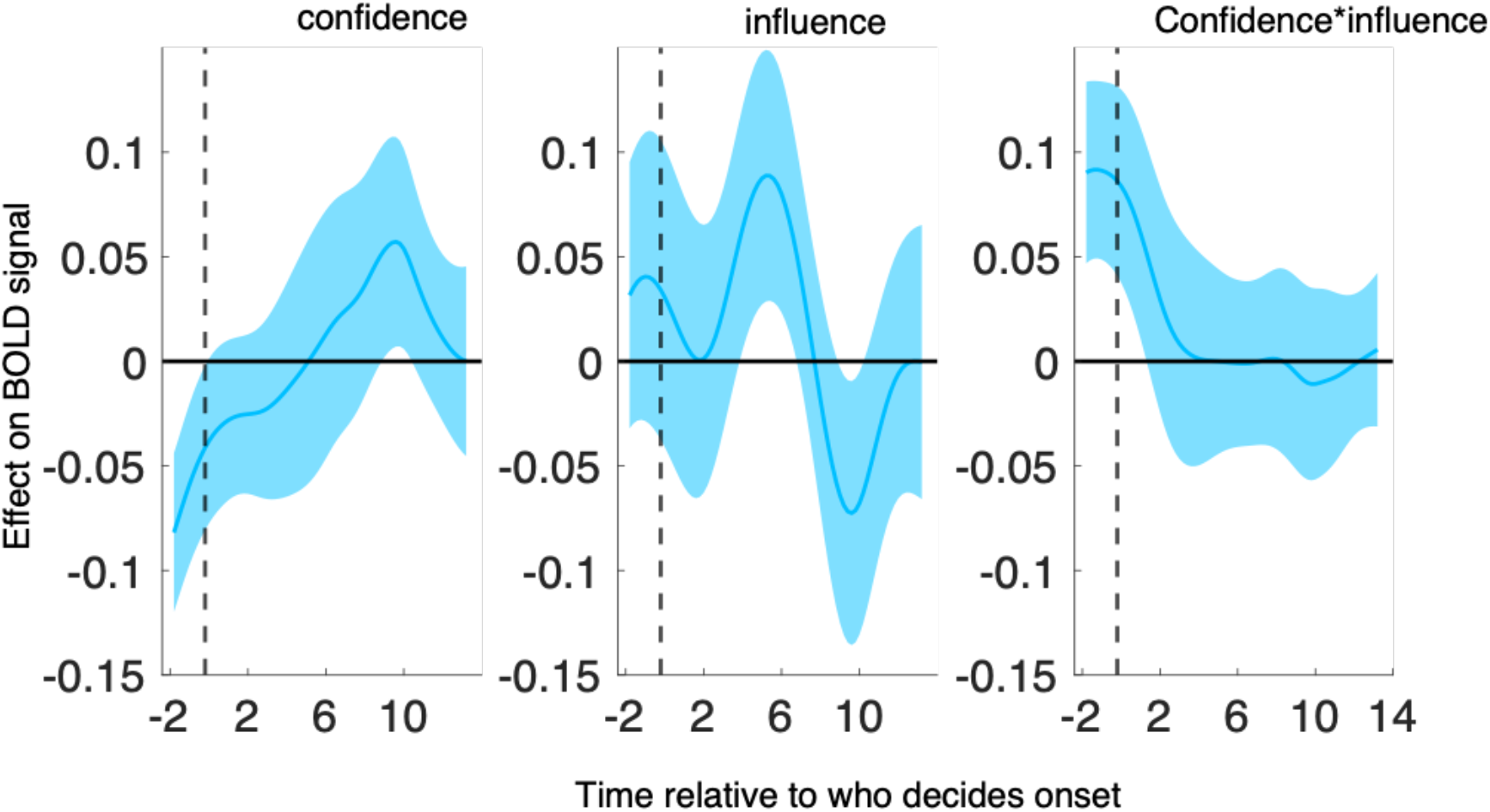
Only for the human condition, we computed response of the dACC to confidence (left), influence (middle), and their interaction (right) when it was announced that the partner would have to announce the revised estimate(See Figure 1). Unlike revise trials where the participants announced the revised decision, in these trials there was no effect of any of the variables on the dACC activity on the observe trials (panel A), the activity.

**Figure S2:**
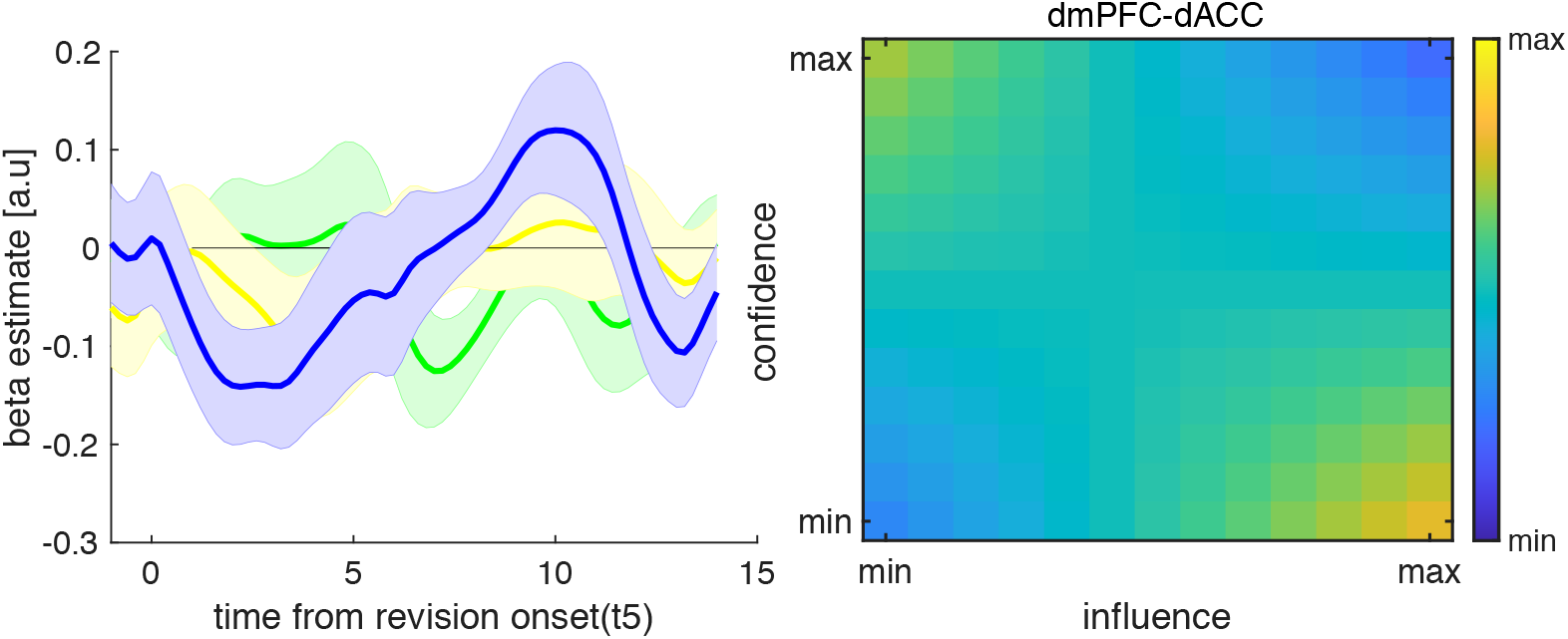
Psychophysiological interaction analysis of ROI activity time courses. Left, Traces are coefficients from a GLM in which we predicted dACC activity from the interaction between dmPFC activity and (1) confidence (yellow), (2) influence (green) and (3) the interaction between confidence and influence (blue) – while controlling for the main effect of each term. Right, Visualisation of dmPFC-dACC connectivity. Hotter colours indicate greater dmPFC-dACC connectivity as a function of variation (in z-score units) in influence (x-axis) and confidence (y-axis). dmPFC-dACC connectivity was estimated using group-level coefficients averaged across a time window from 2 s to 3 s.

### Exploratory whole-brain analysis

Finally, for completeness, we performed an exploratory whole-brain analysis in which we searched for neural correlates of our variables of interest (GLM1). We modelled the task events shown in Figure 1 as separate condition regressors and included separate condition regressors for human and computer blocks (note that some events only happen every other trial). We parametrically modulated a subset of the condition regressors as follows: the periods during which participants made their estimate (visual estimate) and indicated their confidence (confidence rating) were parametrically modulated by confidence; the period during which participants revised their estimate (revision) was parametrically modulated by confidence, the influence that the partner had exerted over participants on the previous trial, their interaction, the degree of revision, the angular distance between the participant’s own and the partner’s first estimate and the angular distance between participants’ revised and the partner’s first estimate; and the period during which participants observed the partner’s revised estimate (revision observation) was parametrically modulated by the influence that participants had exerted over the partner on the current trial.

Here we focus on the whole-brain equivalents of the ROI analyses reported above – see Table S1 for the full set of whole-brain results. At the time of the visual estimate, the whole-brain analysis identified a negative effect of confidence in the precuneus across human and computer conditions (Figure S3, peak coordinates [6 −48 50], k = 129, *t*_*peak*_ ((19) = 4.93, p < .0001). At the time of revision (revision trials only), and consistent with the ROI analysis, the whole-brain analysis identified a positive effect of confidence in a cluster encompassing dACC across human and computer conditions (Figure S4A; peak coordinates [2 −16 42], k = 59, *t*_*peak*_ ((19) = 6.26, p <.0001). Further, in line with previous studies (Cascio et al., 2015), there was a positive effect of amount of revision in ventromedial prefrontal cortex (vmPFC) (Figure S4B peak coordinates [−10 44 −10], k = 26, *p*_*FWE*_ < .05). Finally, we note that the whole-brain analysis did not identify any clusters that tracked influence or its interaction with confidence in the human condition, nor any clusters that tracked the difference between the human and computer conditions at any of the events of interest.

**Figure S3:**
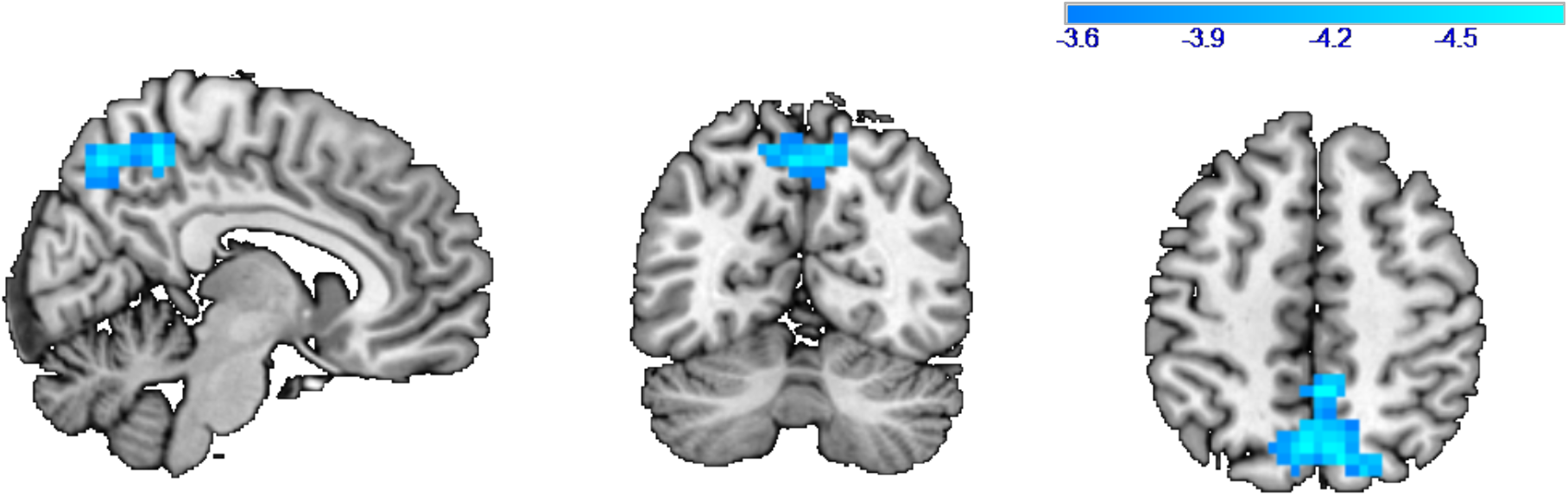
At whole brain, activity of precuneus at the time of first estimate (t2) was significantly negatively modulated by the confidence. Threshold at p<.05, FEW corrected for multiple comparisons, cluster definding threshold p<.0001.

**Figure S4:**
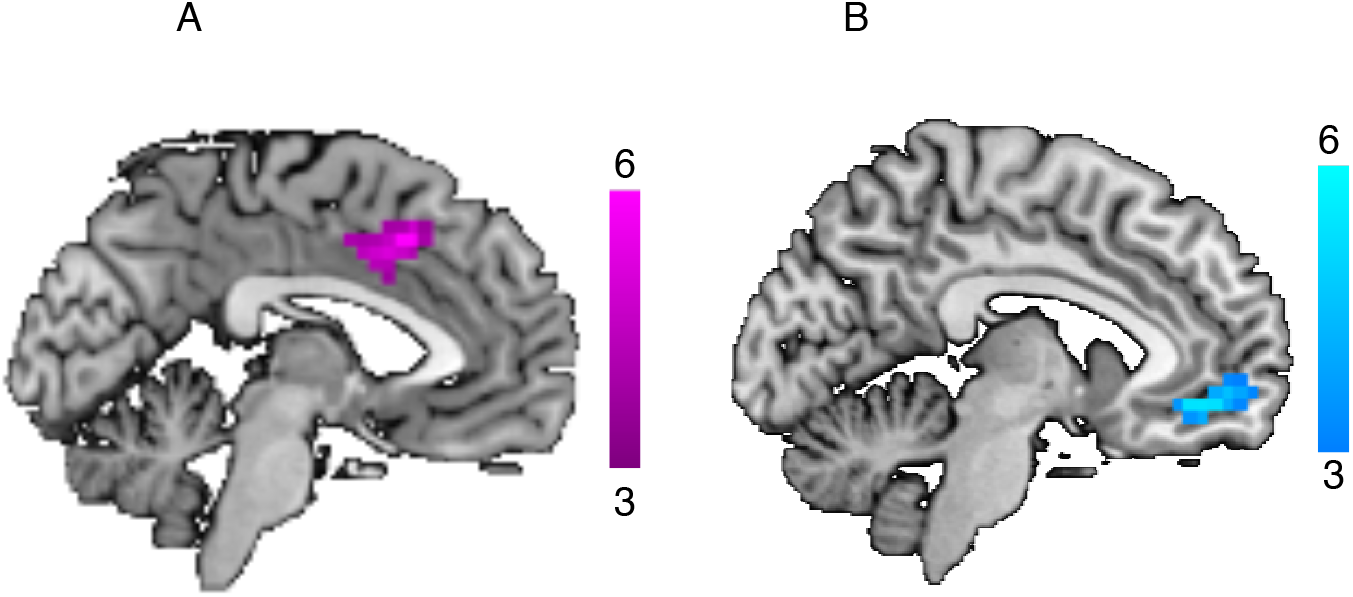
Exploratory whole-brain analysis. Aggregated across both conditions, dACC and vmPFC clusters were modulated by participants’ confidence (left) and their revision toward their partner’s estimate (right) at the time of Revision (t5). Clusters are significant at *P* < 0.05, FWE-corrected for multiple comparisons, with a cluster-defining threshold of P < 0.001, uncorrected.

**Table S1:** different whole brain contrasts which were tested at different event onsets.

| Event | Parametric modulator | contrast |
| --- | --- | --- |
| Estimate | Confidence | Human and computer versus all |
| Confidence rating | Confidence | Human and computer versus all |
| Revision | Confidence | Human and computer versus all |
| Revision | Influence | Human versus all and human versus computer |
| Revision | Influence* confidence | Human versus all and human versus computer |
| Revision | Amount of revision | Human versus all and human versus computer |
| Revision | Angular distance with partner before revision | Human and computer versus all |
| Revision | Angular distance with partner after revision | Human and computer versus all |
| Revision observation | Influence | Human versus all and human versus computer |

### Alternative model for Figure 4 analysis in the main text

For the equivalent fixed-effect model in equation (2) in the main text (where all variables were fixed effect but not random effect) we found a strong negative correlation between the beta weights of influence and the beta weights of the interaction term (Pearson r = −.88, p<.001). This suggests that the weight for influence and interaction are constant across subjects. We, therefore, applied the following modified model:

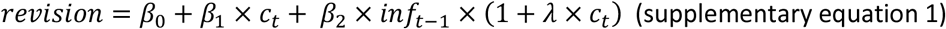

where *λ* is a subject independent parameter while *β*_0_-*β*_2_ can vary across subject. Using a mixed model approach supplementary equation 1 was fitted to the behavioural data for a fixed value of lambda. Lambda was varied on a 1000 point equidistant grid between −2 and 0. This procedure yielded a maximum likelihood estimation of *λ* = −.19.

We therefore also modelled dACC BOLD using this model (supplementary equation 1, replacing revision with dACC BOLD) with the optimised *λ* = −.19.

Consistent with the previous model, in the human condition, confidence had a positive effect on dACC BOLD (Wilcoxon sign-rank test W = 170 P = .01, Figure S5). The effect of influence that was modulated by confidence (*β*_2_ in supplementary equation 1) did not reach significance in this model (Wilcoxon sign-rank test W = 141 P = .09, Figure S5). The absence of an influence effect might be due to greater variability in using normative information among participants. We therefore predicted that the neural effect of influence might be greater in participants who showed a behavioural effect of influence on revision. This means that we expect a correlation between the behavioural effect of influence on revision (*β*_2_ in supplementary equation 1) and the effect of the same regressor on BOLD across participants. To test this hypothesis, we correlated behavioural and neural betas of this regressor across participants. Interestingly, we found that participants with higher behavioural effect of influence on revision, had also greater effect of influence on BOLD (Pearson r = .45, p = .04, Figure S6). Note that, to compute this correlation we controlled for the behavioural and neural betas of confidence using partial correlation. It should be noted that the correlation changes after removing the outlier (Pearson r = .34, p = .18). When we repeated this analysis for the computer partner condition, we found a positive effect of confidence as before, and we did not find an effect of influence on dACC BOLD.

We conclude that both models (modified model and the one used in the main text) lead to the same finding: that confidence and influence have a positive effect on BOLD in dACC, but the effect of influence is negatively modulated by the size of confidence and is restricted to the Human partner condition.

**FigureS5:**
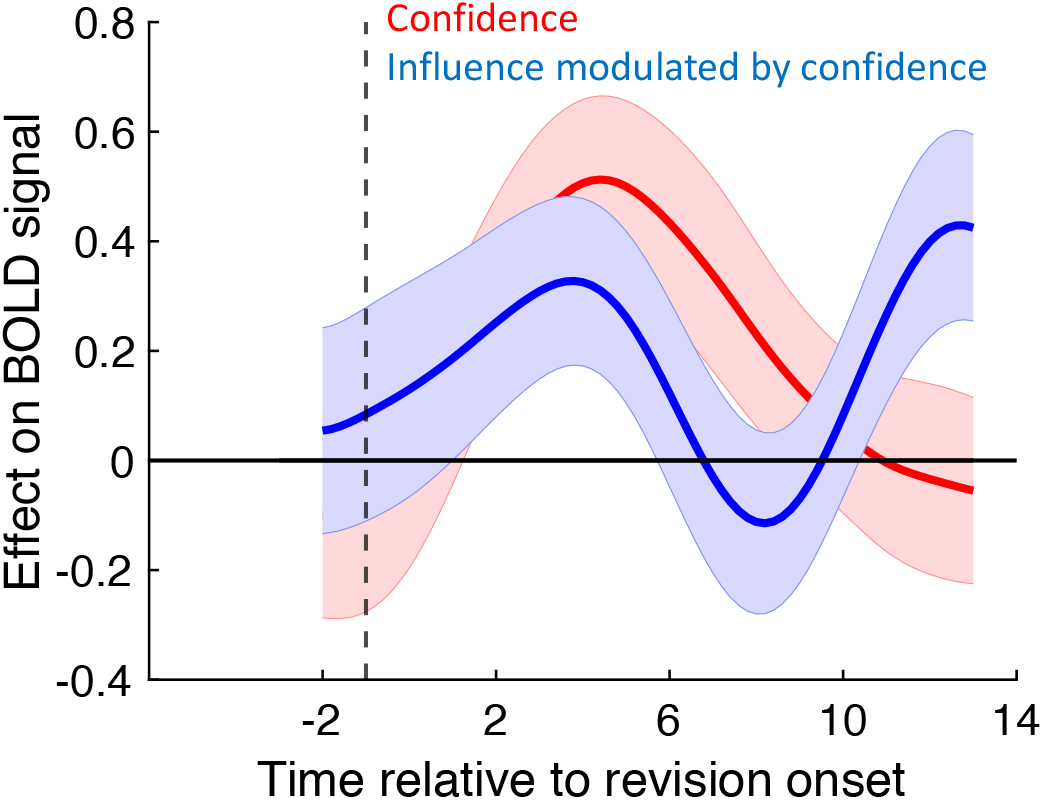
the effect of confidence (red) and influence modulated by confidence using the modified model (blue) on BOLD is depicted.

**Figure S6:**
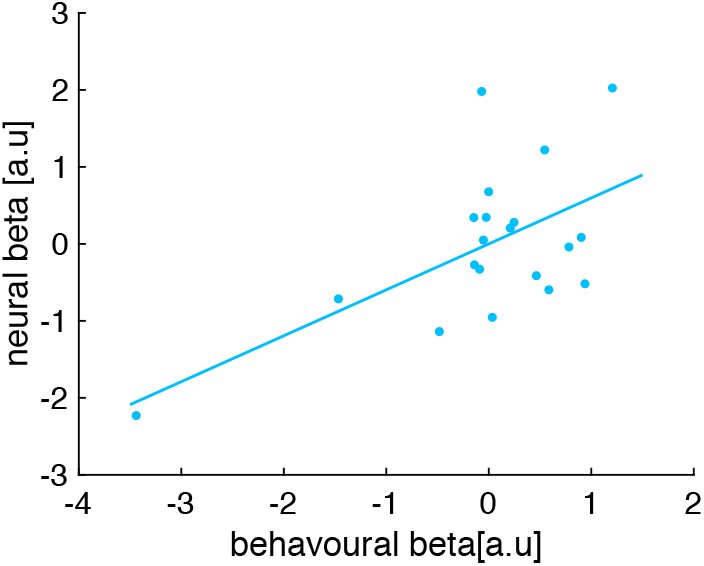
The behavioural (x-axis) and neural (y-axis) effect of influence on dACC BOLD (*β*_2_ in supplementary equation 1) is depicted for each participant.

